# Rhizosphere and detritusphere habitats modulate expression of soil N-cycling genes during plant development

**DOI:** 10.1101/2023.03.24.534069

**Authors:** Ella T. Sieradzki, Erin E. Nuccio, Jennifer Pett-Ridge, Mary K. Firestone

**Author notes:** Current address: Laboratoire Ampère, École centrale de Lyon, France (Ella T. Sieradzki).

## Abstract

Interactions between plant roots and rhizosphere bacteria mediate nitrogen (N)-cycling processes and create habitats rich in low molecular weight (growing roots, rhizosphere) and complex organic molecules (decaying root litter, detritusphere) compared to bulk soil. Microbial N-cycling is regulated by a diverse suite of genes from many interconnected metabolic pathways; but most studies of soil N-cycling gene expression have focused on single pathways. Currently, we lack a comprehensive understanding of the interplay between soil N-cycling gene regulation, spatial habitat and time. Here we present an analysis of a replicated time series of soil metatranscriptomes; we followed multiple N transformations in four soil habitats (rhizosphere, detritusphere, mixed rhizo-/detriusphere, bulk soil) over a period of active root growth for the annual grass, *Avena fatua*. The presence of root litter and living roots significantly altered the trajectory of N-cycling gene expression. Across soil habitats, the most highly expressed N-transformation genes were related to extracellular proteases, ammonium assimilation into microbial biomass via glutamate synthase, and ammonium oxidation. Upregulation of bacterial assimilatory nitrate reduction in the rhizosphere suggests that rhizosphere bacteria were actively competing with roots for nitrate. Simultaneously, bacterial ammonium assimilatory pathways were upregulated in both rhizosphere and detritusphere soil, which could have limited N availability to plants. The detritusphere supported dissimilatory processes DNRA and denitrification. Expression of ammonium oxidation genes was almost exclusively performed by three phylotypes of *Thaumarchaeota* and was upregulated in unamended bulk soil. Unidirectional ammonium assimilation and its regulatory genes (glutamine synthetase/glutamate synthase, or GS/GOGAT) were upregulated in soil surrounding relatively young roots and more highly decayed root litter, suggesting N may have been limiting in these habitats (the GS/GOGAT pathway is known to be activated under low N availability). We did not detect expression of N-fixation or anammox genes. Our comprehensive metatranscriptomic time-series of organic and inorganic N-cycling in rhizosphere, detritusphere, and bulk soil, indicates that differences in C and inorganic N availability control contemporaneous transcription of N-cycling pathways in soil microhabitats that exist in close spatial proximity.

**Contribution to the field:** Plant roots modulate microbial nitrogen cycling by regulating the supply of root-derived carbon and nitrogen uptake. These differences in resource availability cause distinct micro-habitats to develop: soil near living roots (rhizosphere), decaying roots (detritusphere), near both (rhizo/detritusphere), or outside the direct influence of roots (bulk). While many genes control the microbial processes involved in the nitrogen cycle, most research has focused on single genes and pathways, neglecting the interactive effects these pathways have on each other. The processes controlled by these pathways determine consumption and production of N by soil microorganisms. We followed the expression of N-cycling genes in the primary four soil microhabitats over a period of active root growth for an annual grass. We found that the presence of root litter and living roots significantly altered gene expression involving in multiple nitrogen pathways. We also found populations with genes for multiple pathways, where expression was likely shaped by available forms of carbon and by competition with plants for inorganic nitrogen. Phylogenetic differences in spatial and temporal expression of the soil microbial N-pathway genes ultimately regulate N-availability to plants.

## Introduction

Nitrogen (N) is a key limiting nutrient for plant growth, since in soil, N mostly occurs in organic forms within large, complex molecules that are unavailable to plants. Soil microorganisms have suites of extracellular enzymes capable of degrading these molecules, freeing small organic and inorganic N to the surrounding soil and benefitting nearby plants. Other bacteria and archaea transform N to nitrate—an inorganic form that is more available to many plants. Meanwhile, microorganisms also take up inorganic N for a variety of growth-supporting assimilatory and dissimilatory metabolic processes. This can lead to some competition for N between soil bacteria and plants. All of these processes likely occur simultaneously in soil, but we have an incomplete understanding of how they vary with time and in distinct soil habitats.

Soils contain numerous microhabitats with heterogeneous distributions of substrates and resources (Sokol et al., 2022). The influence of growing roots can dominate (rhizosphere) or dead root litter may dominate (detritusphere); alternatively these two habitats may co-occur when live roots regrow into previously colonized areas (rhizo/detritusphere). In previous analyses, we have shown that distinct bacterial guilds operate in rhizosphere, detritusphere, and combined rhizosphere-detritusphere soil habitats (Nuccio et al., 2020). The spatial organization of these soil habitats may be particularly important for nutrient transfers that create distinct N microhabitats that enable or limit N-cycling. For example: (1) in the rhizosphere, exudate-driven blooms of microbial growth are followed by predation that liberates the nutrient capital held in bacterial, fungal, and metazoan bodies, this redistribution of microbial biomass causes conversion to lower molecular weight compounds (*e.g*., amino acids) that are deaminated to NH_4_^+^ by diverse bacteria and fungi (Clarholm, 1985; Koller et al., 2013); (2) in the detritusphere, organic N in root litter (e.g., lignoproteins (Myrold, 2021)) is mineralized to make inorganic N available soil microbes, and also serves as a carbon source for soil microbes (Pepe-Ranney et al., 2016), creating a demand for N to maintain cellular stoichiometry; (3) in bulk soils, microbes may express more macromolecule degradation genes relative to more resource rich sites near roots, where degradation enzymes for low molecular weight compounds are more prevalent (Shi et al., 2018).

Rates of microbial N transformations in soil can be significantly impacted by the presence of both growing and decaying roots. Gross N-mineralization, immobilization, and nitrification rates vary as a function of proximity to plant roots, root age and concentration of organic material in grassland soil (Personeni and Loiseau, 2005; Herman et al., 2006). Roots can significantly deplete ammonium in the rhizosphere within days of root introduction into sterile soil (Trofymow et al., 1987). However, in wild soils with intact microbial communities, rhizosphere-associated bacteria can transiently compete with plants (e.g., *A. barbata* roots) for inorganic nitrogen (Jackson et al., 1989a). As part of this competition, plants can inhibit rhizosphere nitrogen transformations such as biological nitrification inhibition (BNI) via production of certain root exudate compounds (Nardi et al., 2020). Plants have also been shown to inhibit both nitrification and denitrification processes in the rhizosphere—both processes lead to loss of N from the plant-available pool (Moreau et al., 2019)—and litter amendment has been shown to increase denitrification (Che et al., 2018). Understanding the N-cycling tradeoffs between different soil habitats is key for our ability to better model and predict the controls of soil N cycling.

Soil metatranscriptome analysis is a powerful tool for linking nitrogen cycling genes to specific microbial populations, and correlating transcription levels to environmental conditions (Nicol et al., 2008; Liu et al., 2010; Placella and Firestone, 2013; Nuccio et al., 2020; Sieradzki et al., 2020; Tosi et al., 2020). In comparison, qPCR can target only single genes, and primer biases may leave some taxa undetected. A recent metatranscriptomics study demonstrated a short-term coupling of nitrogen cycling pathways and availability of simple carbons simulating priming by root exudates (Chuckran et al., 2021). Otherwise, there is little knowledge on the contemporaneous feedback through gene expression between carbon availability and nitrogen cycling in naturally complex soil habitats and over annual plant-relevant time frames (Yergeau et al., 2014, 2018; Žifčáková et al., 2016).

Here, we used soil metatranscriptomics analysis to build a comprehensive time-resolved representation of both organic and inorganic nitrogen transformations in an annual grassland soil. We characterized a period of active root growth, as well as root decomposition, of wild oat grass (*Avena fatua*), a common species in Mediterranean grasslands. We hypothesized that different N cycle pathways would be prevalent in different soil microhabitats that vary in available C and N resources. We analyzed 48 metatranscriptomes from rhizosphere and bulk soil collected over a 3-week time series in the presence or absence of root litter; this allowed us to identify the dominant N-cycling pathways associated with each habitat and untangle the effects of carbon supply by live or dead roots from competition with the plant for N. We previously analyzed a different subset of this data to assess expression of genes coding for carbohydrate degradation (Nuccio et al., 2020) and organic nitrogen degrading enzymes (Sieradzki et al., 2020). While these prior studies provide important context, our analysis here is unique in its comprehensive exploration of how N transformation gene expression is influenced by the distinct environmental conditions of different soil habitats. The use of metatranscriptomics (as opposed to qPCR) also facilitated identification of specific microbial populations involved in each pathway, and helped us test for multiple N transforming pathways expressed within populations found in mutiple soil habitats. Finally, we measured changes in N-pathway gene expression over time as the rhizosphere aged and as root litter was degraded.

## Methods

### Experimental design, sample collection and sequencing

The experimental procedures and initial data processing for our 48 soil metatranscriptome dataset are described in detail by (Nuccio et al., 2020). Briefly, *Avena fatua* was grown in a fine loam Alfisol complex (Ultic Haploxeralf mixed with a Mollic Palexeralf) from the Hopland Research and Extension Center (Hopland, CA); pH 5.6, 2% total C) in microcosms with a sidecar with transparent walls. After six weeks, roughly halfway through the plant life span, the solid divider to the sidecar was removed and replaced with a slotted divider so that roots could grow into the sidecar. Root growth was marked on the sidecar wall. All sidecars contained bulk soil bags that were inaccessible to roots. Half of the bulk soil bags as well as the soil in half of the sidecars were amended with dried *A. fatua* root detritus (litter C:N = 13.4). Once the sidecar was opened, paired rhizosphere and bulk soil triplicates were harvested destructively after 3, 6, 12 and 22 days from amended and unamended microcosms for a total of 48 samples. Lifeguard Soil Preservation Reagent (MoBio) was added to 1g subsample of the harvested soil. The supernatant and roots were removed, and the samples were stored at −80°C.

DNA/RNA co-extraction was performed with phenol-chloroform. DNA and RNA were separated with an AllPrep kit (Qiagen) and RNA was DNase treated (TURBO DNase, Thermo-Fisher Scientific). Ribosomal RNA was depleted (RiboZero, Illumina) and the mRNA was retrotranscribed. cDNA was sequenced with an Illumina HiSeq 2000 2×150 (TruSeq SBS v3) protocol at the Joint Genome Institute. In addition, the V4 hypervariable region of the 16S gene (primers 515F and 806R) was amplified from the extracted DNA and sequenced via Illumina MiSeq v3 2×300. Amplicons were analyzed by the Joint Genome Institute. Operational taxonomic units (OTUs) were clustered at 97% identity with USEARCH (Edgar, 2010) and taxonomy was assigned at 95% id by RDP (Wang et al., 2007).

### Expression of genes from a curated collection of Hopland-soil genomes

As described previously (Nuccio et al., 2020), raw reads were quality-trimmed (Q20) and rRNA and tDNA were removed. Library size, evenness, richness and Shannon diversity were comparable between experimental groups, with a mean library size of 43 M paired end reads. Reads were mapped at 80% identity with BBsplit (Bushnell, 2014; Nuccio et al., 2020) against a dereplicated reference set of 282 Hopland CA soil genomes including stable isotope probing metagenome assembled genomes (SIP-MAGs) (Starr et al., 2018), metagenomic assembled genomes (MAGs, NCBI PRJNA517182), soil isolates (Starr et al., 2018; Zhalnina et al., 2018) and single amplified genomes (SAGs) (Nuccio et al., 2020). A mean of 12.3% of the reads per sample mapped unambiguously to the reference set.

Prodigal (Hyatt et al., 2010; Nuccio et al., 2020) was used to predict open reading frames (ORFs) in the genomes. The ORFs were annotated using KEGG (Kanehisa and Goto, 2000) and ggKbase (http://ggkbase.berkeley.edu). Gene counts were calculated with R package Rsubread function featureCounts (Liao et al., 2014). Three samples were removed from further analysis due to an exceptionally low number of features: H3_Rhizo_NoLitter_39, H1_Rhizo_Litter_2 and H2_Rhizo_Litter_9.

All features and their metatranscriptomic read recruitment were analyzed with the R package DESeq2 (Liao et al., 2014; Love et al., 2014) requiring an adjusted p-value < 0.05. Ordination and visualization were conducted using R package ggplot2 (Wickham, 2016) and vegan (Oksanen et al., 2008; Wickham, 2016); the vegan function Adonis was used to detect significant factors affecting expression of nitrogen cycling genes.

### Expression of nitrogen cycling genes identified in assembled metatranscriptomes

After quality control, the metatranscriptomes were assembled into contigs within samples. Only contigs larger than 200 bp were clustered at 99% ID with cd-hit-est (Huang et al., 2010). Prodigal (Huang et al., 2010; Hyatt et al., 2010) was used to predict ORFs from the cluster representatives, and KEGG models (Kanehisa and Goto, 2000) were used to locate nitrogen cycling genes based on the KEGG nitrogen cycle module (see Supplemental Table S1 for a list of KEGG profiles, gene lengths and thresholds used). Some ORFs were identified by several HMMs due to homology (e.g. nitrate reductases *nxrA*, *narG* and *napA*). In those cases, the highest scoring (by bitscore) hit from all HMMs was selected. Extracellular proteases were called by reciprocal blast to proteases from the MEROPS database that contained signal peptides (Rawlings et al., 2016; Nguyen et al., 2019). Reads were mapped to nitrogen cycling ORFs at a minimum identity 95% and minimum breadth 75% using bbmap (Bushnell, 2014; Nuccio et al., 2020). Normalization to sequencing depth was done using DESeq2, and when comparing between different genes expression was also divided by the relevant mean assembled gene length to control for gene-length biases in recruitment. Heat maps were generated in R using gplots (Warnes et al., 2015).

Phylogeny of the ammonia monooxygenase subunit A (*amoA*) was determined by aligning *amoA* assembled transcripts to curated full amoA gene sequences from RefSeq (Jun 1, 2019) using mafft-linsi (sensitive mode) (Katoh and Standley, 2013) and removing positions represented by less than 50% of the aligned sequences with GBlocks version 0.91 (Dereeper et al., 2008) with parameters −b3=50 −b4=5 −b5=h. RAxML version 8.2.12 (Kozlov et al., 2019) with parameters set to matrix LG and seed=13 was used to build a Newick tree which was visualized in iTOL (Letunic and Bork, 2019). The phylogeny of *nirK* transcripts was determined in the same manner. The aggregated coverage of *amoA*/*nirK* variants from all samples was superimposed onto the tree in iTOL. Additionally, since nitrate reductases that participate in multiple pathways are homologous, we wanted to make sure they were correctly identified by the HMMs. Therefore, we placed the assembled sequences on a protein reference tree (Matheus Carnevali et al., 2019) and corrected the annotations according to tree clustering.

Normalized expression per gene per time point was compared between groups using ANOVA and Tukey HSD tests, and p-values were corrected for multiple comparisons (time, location, amendment and all interactions between them). Upregulation was determined as log2-fold values calculated by DESeq with corrected *p* values < 0.05.

## Results

To assess whether expression of organic and inorganic nitrogen cycling genes was affected by the presence of root litter in the rhizosphere or in nearby bulk soil, we tested for significant differences in N transformation gene expression over a three-week time series (3, 6, 12, 22 days). The PCoA based on Bray-Curtis dissimilarity of expression of N cycling genes normalized to sequencing depth revealed clustering by location (rhizosphere/bulk soil) throughout the whole time-series; the last time point (T4; 22 days) notably distinct from the others (fig. 1A). Analysis of the first three time points only (3, 6, and 12 days) revealed an additional clustering by litter amendment (fig. 1B). In both analyses, the first two principal components explained 40% of the variation. PERMANOVA analysis using all time points indicated a significant effect of location (rhizosphere vs. bulk soil), time and litter amendment (p-value = 10^-4^) (see sup. table 1 for all values). The combined R^2^ of all significant factors was 63%. When we conducted the same analysis but excluded the last time point (T4), the effects of single factors remained significant (p-value = 10^-4^). The combined R^2^ of all significant factors was 61%, and the variability explained by time decreased from 28% with T4 to 10% without T4, whereas the variability explained by location and amendment increased by 5-8% each.

**Figure 1:**
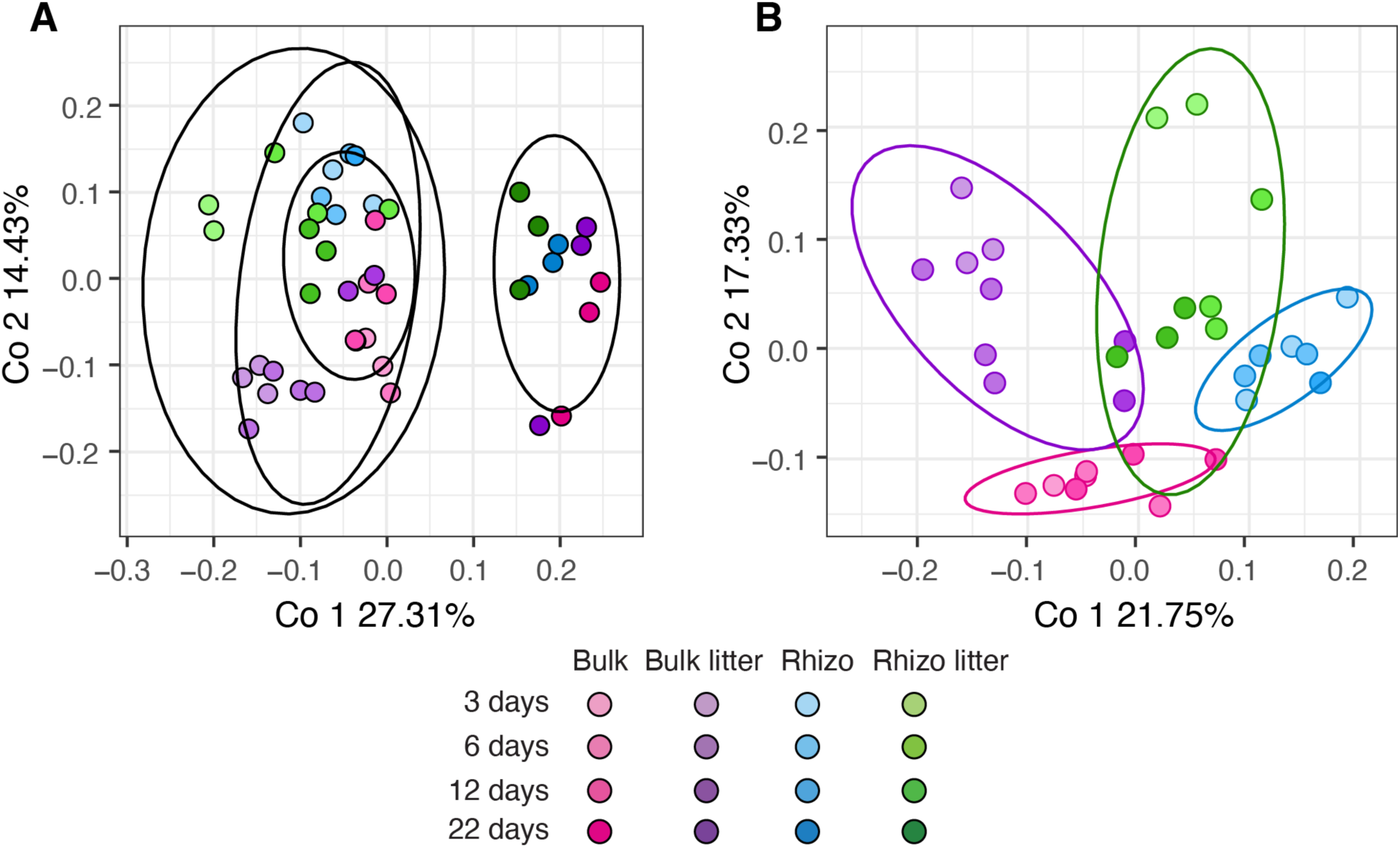
Principal coordinates analysis (PCoA) of nitrogen cycling gene expression. Influence of living roots and root detritus on soil microbial gene expression during 3 weeks of *Avena fatua* root growth (harvests at 3, 6, 12, 22 days) in a CA annual grassland soil. Input data are Bray-Curtis dissimilarity of expression of N cycling genes normalized to sequencing depth, determined by mapping reads to ORFs assembled from metatranscriptomes followed by log ratio transformation, in (A) all time points with the last time point circled and (B) excluding the last time point (22 days). Groups are indicated in the ellipses: B=Bulk, BL=Bulk Litter, R=Rhizosphere, RL=Rhizosphere Litter. n=3 for each habitat and timepoint. Ellipses represent a 90% confidence interval of the ordination coordinates (calculated by stat_ellipse).

### Expression of nitrogen cycling genes by pathway

To identify the dominant N-cycling pathways in four soil microhabitats (rhizosphere, detritusphere, rhizo/detritusphere, bulk soil), we compared differential expression of each nitrogen cycling gene in the presence or absence of roots and litter amendment. Expression and upregulation trends of N cycling genes (defined in Table S1) are shown in fig. 2. Several genes involved in assimilatory pathways were significantly upregulated in the rhizosphere: assimilatory nitrate reduction genes *nasA*, *NR* and *nirA*, glutamate dehydrogenase *gdh2* and GS/GOGAT genes *glt1* and *gltB* (fig. 2). The first step of GS/GOGAT, *glnA*, was upregulated both in the rhizosphere and in the presence of litter (fig. 2), and the GS/GOGAT regulatory proteins *glnD* and *glnG*, triggered under N limitation, were upregulated in the rhizosphere, whereas *glnB* was upregulated in the presence of litter (sup. fig. 1 B,C). In the detritusphere, the presence of litter also triggered macromolecular organic nitrogen degradation (extracellular protease and chitinase), as well as dissimilatory pathways; dissimilatory nitrate reduction (DNRA) and denitrification (fig. 2). However, not all genes in these pathways were upregulated and some were not detected at all (*nirS* and *norBC* in denitrification), whereas others were not significantly upregulated (*nirK*, *narGHI*, *napB*) (fig. 2). Finally, nitrification, DNRA and the first step of denitrification were upregulated in bulk soil compared to the rhizosphere (sup. fig. S1).

**Figure 2:**
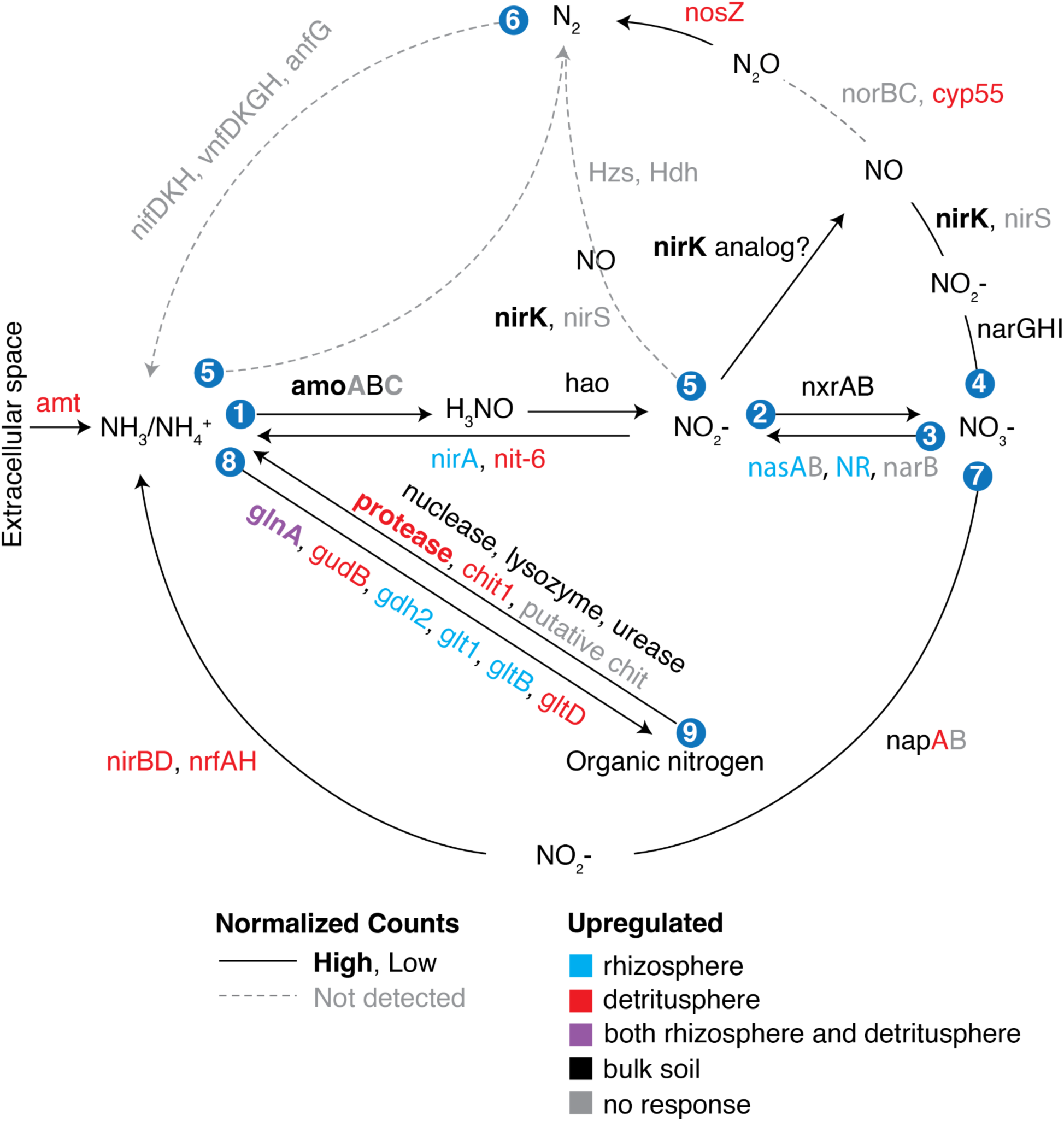
Expression level and upregulation by soil habitat of nitrogen cycling genes from assembled transcripts. Bold font indicates high expression levels (mean counts in each time point >5 after normalization to gene length and sequencing depth). Significant upregulation was determined by ANOVA and Tukey post hoc test (adjusted p<0.05). Numbers denote pathways: 1=Ammonia oxidation, 2=Nitrite oxidation, 3=Assimilatory nitrate reduction, 4=Denitrification, 5=Anammox, 6=Nitrogen fixation, 7=DNRA, 8=Ammonium assimilation (GDH and GS/GOGAT), 9=Macromolecular N mineralization, amt = Ammonium transporter. The functional role of each enzyme is explained in Table S2. The nirK analog is an archaeal enzyme which participates in archaeal nitrification, but unlike its bacterial analog, it does not generate nitrate (Stein, 2019).

Several genes were particularly highly expressed in our dataset (e.g., ammonia monooxygenase subunits *amoA* and *amoC*, and nitrate reductase *nirK* (sup. fig. S1 A,B)) but did not show clear patterns of differential expression by either soil habitat treatment or time. We did not detect any genes involved in nitrogen fixation or anaerobic ammonia oxidation. Expression of fungal N cycle genes such as *cyp55* (nitric oxide to nitrous oxide) and *nit-6* (nitrite to ammonium) was very low (sup. fig. 1C).

### Archaeal nitrification

Two of the most highly expressed genes we identified were subunits of ammonium monooxygenase, the first enzyme in the nitrification pathway. Nitrification genes were not affected by the presence of litter, but *nirK*, *amoB*, *hao*, *nxrA* and *nxrB* were significantly upregulated in bulk soil compared to rhizosphere soil (ANOVA: p<0.05, Tukey HSD test: adjusted p<0.05) (sup. table S2).

Ammonium oxidation can be performed either by ammonium oxidizing archaea (AOA) or ammonium oxidizing bacteria (AOB). The *amoA* subunit of ammonia monooxygenase is commonly used as a taxonomic marker to differentiate between AOA and AOB. To determine which taxa were responsible for the high expression of ammonia oxidation genes, we placed the 13 *amoA* ORFs identified in assembled transcripts into a phylogenetic tree with full-length *amoA* reference sequences from the RefSeq database (fig. 3A). The cumulative coverage per *amoA* variant in all samples was superimposed onto the tree. Read recruitment to AOA was orders of magnitude higher than to AOB, accounting for 98% ± 0.007% (N=16) of the reads mapped to *amoA* per sample. While there were some ORFs placed into comammox clade B, their expression was negligible compared to archaeal *amoA* (fig. 3A).

**Figure 3:**
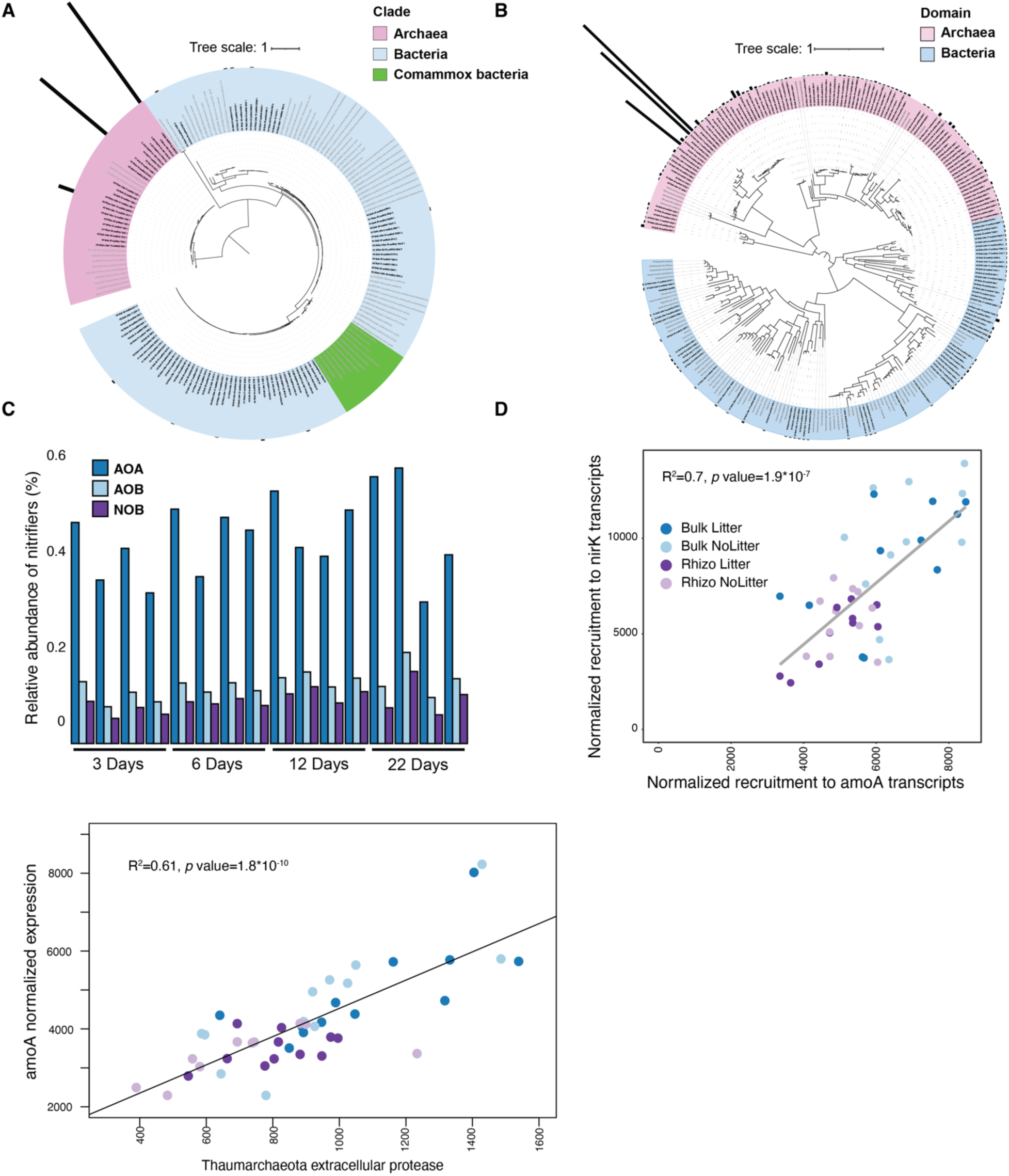
Phylogenetic determination of nitrifiers by *amoA*, *nirK* and *16S-rRNA*. Phylogeny of assembled ORFs (black) (A) *amoA* and (B) *nirK* placed into phylogenetic trees containing curated reference sequences from RefSeq (grey). Colors represent phylogenetic clades. Bars around the trees represent cumulative coverage from all samples to specific assembled transcripts. (C) Relative abundance of AOA, AOB and nitrite oxidizing bacteria (NOB) of the entire microbial community by *16S-rRNA* gene abundance. (D) Positive correlation between expression of *amoA* and *nirK* transcripts by location and amendment (R^2^=0.7). (E) Positive correlation between expression of *amoA* and *Thaumarchaeota* extracellular protease transcripts by location and amendment (R^2^=0.61).

The expression of hydroxylamine oxidoreductase (*hao*), the next step in nitrification after ammonium oxidation, was extremely low across all samples. Similarly, expression of nitrite oxidase (*nxrAB*) was very low. However, the expression of *nirK* was very high (fig. 2). A phylogenetic analysis placed 96 out of 158 *nirK* assembled ORFs in the archaeal clade and, as in the case of *amoA*, revealed that the archaeal variants recruited more than 95% ± 0.07% (N=16) of the reads that mapped to *nirK* (fig. 3B). Three archaeal phylotypes dominated *nirK* expression, with 76% ± 0.07% of all reads mapped to *nirK*.

To determine whether the high expression of archaeal nitrification genes can be explained by abundance of AOA compared to AOB, we searched our *16S rRNA* gene amplicons (Nuccio et al., 2020) for known AOA and AOB. The relative abundance of AOA *16S rRNA* genes was 2- to 5-fold higher than for AOB, and the relative abundance of AOB was consistently higher than that of nitrite-oxidizing bacteria (NOB) (fig. 3C). The aggregate expression of *amoA* and *nirK* was well correlated with a ratio of *nirK*:amoA = 1.6 (R^2^=0.7, *p*<0.0001) (fig. 3D). In addition, the aggregated expression of amoA was also highly correlated to that of Thaumarchaeal extracellular proteases (Sieradzki et al., 2020) (adjusted R^2^=0.61, *p*<0.0001).

### Ammonium assimilation and transport pathways

The most highly expressed ammonia acquisition pathway was glutamate synthase (GS/GOGAT), which can be activated by a cascade of four enzymes under nitrogen limitation (fig. 4A) (MacNeil et al., 1982; Bueno et al., 1985; Leigh and Dodsworth, 2007). As opposed to single-gene dependent transporters (*amt*) and glutamate dehydrogenase (GDH), GS/GOGAT is a two-step pathway: the first step is encoded by the gene *glnA*, and the second step either by *glt1, Glu* or the heterodimer *gltBD*. Both ammonium transporters (sup. fig. 1) and GS/GOGAT (*glnA*, fig. 4B) were significantly upregulated in detritusphere microhabitats within the young rhizosphere (Rhizo Litter, 3 days) and bulk soil at the final timepoint (Bulk Litter, 22 days) (fig. 4b). *glnA* was also significantly upregulated in the early rhizosphere compared to the bulk control (Rhizo, fig 4B), and three out of four detected proteins involved in GS/GOGAT were also upregulated in the rhizosphere after 3, 6 and 12 days (sup. fig. S3). We found that expression of *gltBD* was generally much higher than that of *glt1*, whereas *Glu* was not detected (sup. fig. 1). Expression of an additional ammonium assimilation pathway, glutamate dehydrogenase (GDH), was low (sup. fig. S1). GDH consists of a single step performed by one of two enzymes: *gdh2* or *gudB*. *Gdh2* was upregulated in the rhizosphere, whereas *gudB* was upregulated in the presence of litter (sup. fig. S1, sup. table S2).

**Figure 4:**
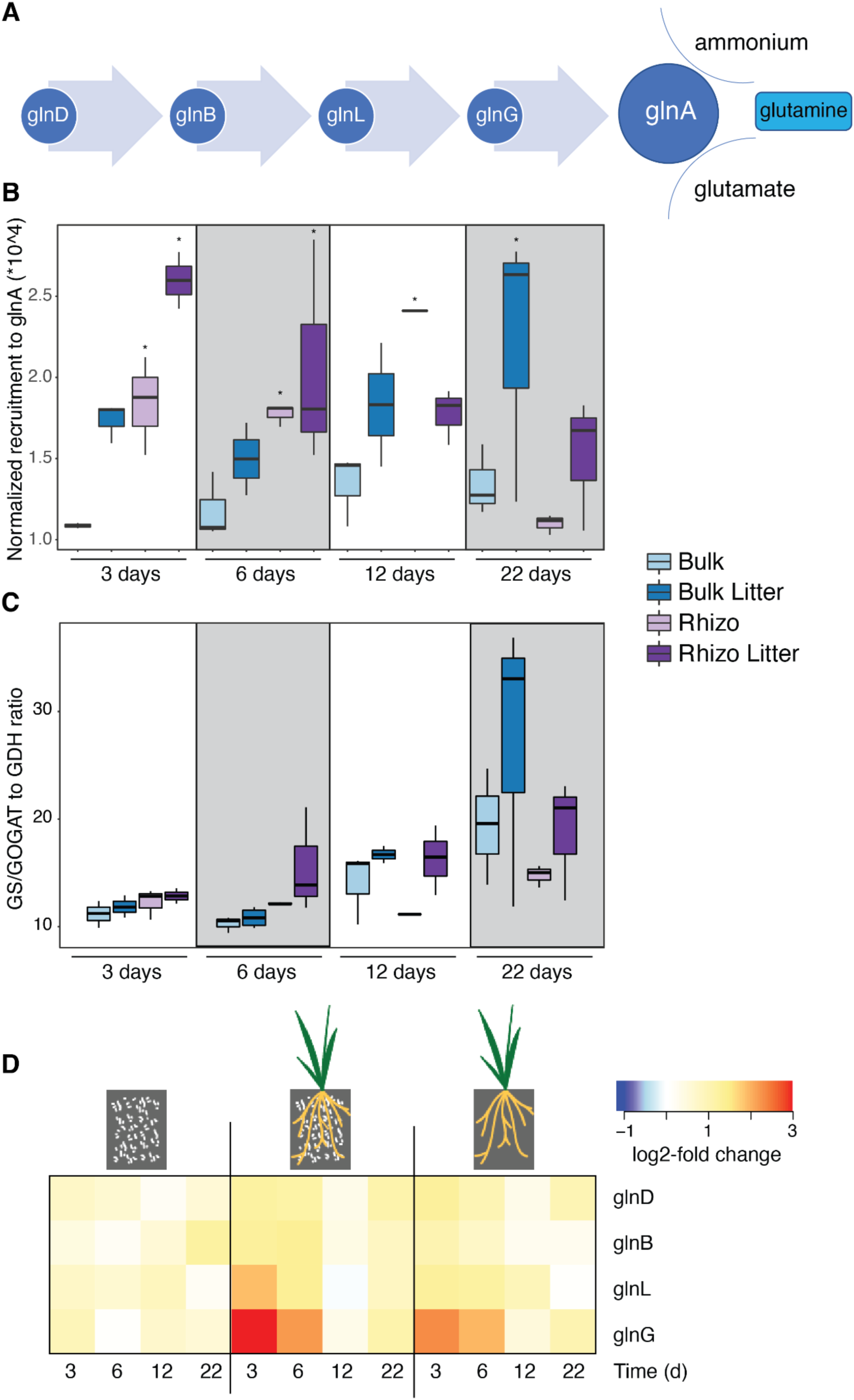
Expression of ammonium assimilation pathways and their regulatory genes. (A) Simplified conceptual representation of *glnA* activation via the regulatory cascade (B) Normalized expression of *glnA* in amended/unamended rhizosphere and bulk soil. Asterisks denote a significant increase compared to the same litter treatment in bulk soil (e.g., unamended rhizosphere vs. unamended bulk) (C) ratio of GS/GOGAT to GDH (*glnA*:*gudB)* expression in the metatranscriptomes. T. (D) Mean differential expression compared to unamended bulk soil of regulatory genes that activate GS/GOGAT under N limitation as well as *glnA*.

In theory, there is an advantage to using GS/GOGAT for ammonia assimilation over GDH under N limitation, as GS/GOGAT in a one-way pathway, whereas GDH can also catalyze deamination of glutamate which might lead to loss of ammonium via diffusion. Expression of GS/GOGAT was an order of magnitude higher than that of GDH under all experimental conditions (fig. 4C). The ratio of expression (GS/GOGAT to GDH) significantly increased over time (Permanova ratio~group*time, p(time)=0.002). In the young rhizosphere the ratio appeared to be higher in the rhizosphere compared to bulk soil, whereas in the mature rhizosphere, litter seemed to trigger a higher ratio (fig. 4D). Expression of the GS/GOGAT regulatory proteins *glnD, glnG and glnL* was significantly upregulated in the rhizosphere compared to bulk soil, particularly in the young rhizosphere (Table S3). Most regulatory genes were also upregulated in the detritusphere compared to unamended soil at the last time point.

## Discussion

In this study we used metatranscriptomic gene expression (% of reads mapped to a gene) and upregulation (fold change of expression in a habitat compared to unamended bulk soil) to identify contemporaneous N transformations in the main soil habitats: rhizosphere (near live roots), detritusphere (near dead roots), rhizo-detritusphere (live and dead roots) and bulk soil (neither). We found that expression of genes coding for N-cycling processes varied between soil habitats as well as over time. The use of metatranscriptomic analyses allowed us to track multiple N cycling pathways occurring at the same time. Moreover, as N pools in this grassland soil are relatively constant but fluxes are high, it has been suggested that studying N cycling parameters may be a better proxy than pools (Sudderth et al., 2012).

Habitat and temporal patterns of N cycling gene expression were interrelated. Effects of habitat on gene expression were detected at the first three sampling times, but not after 22 days. At this time point, the soil was much drier due to plant transpiration (Nuccio et al., 2020). This decrease in soil moisture likely affected rates of substrate diffusion as well as microbial mobility, leading to changes in the environmental milieu experienced by the microbes. Removing the last time point from analyses, revealed strong effects of roots and litter as well as the interaction between the two, implying that rhizosphere bacteria respond to litter amendment differently than bulk soil bacteria. At 22 days, the roots present in the rhizosphere treatment may have begun senescence and degradation processes more similar to that of root litter. Moreover, root exudation decreases as the root senesces, thus likely reducing the effect of simple carbon sources originating from exudation as well as driving usage of litter, which is a less labile C source (Shi et al., 2018).

Several N cycling pathway genes were differentially expressed in different soil habitats. In the rhizosphere we identified upregulation of assimilatory nitrate reduction and ammonium assimilation through glutamate dehydrogenase (GDH; *gdh2*) and glutamine synthetase GS/GOGAT; *glnA, glt1, gltB*). An additional GDH enzyme, *gudB*, was upregulated in the litter-amended rhizosphere. Three of four regulatory enzymes that activate GS/GOGAT under N limitation were also upregulated in the rhizosphere, and the fourth was upregulated in the litter-amended rhizosphere. The upregulation of these pathways likely indicate that the rhizosphere microbial community experiences inorganic N limitation. N limitation can stem from competition over inorganic N with the plant (Jackson et al., 1989b; Schimel et al., 1989), as well as from a higher demand for N as microbial density is ca. 10X higher in the rhizosphere compared to bulk soil (Pett-Ridge et al., 2021). *Avena* plants have been shown to be better competitors than their rhizosphere community for nitrate, potentially explaining why rhizosphere microorganisms would upregulate nitrate reduction to ammonium.

Pathways upregulated in the detritusphere compared to bulk soil were dissimilatory nitrate reduction (DNRA), denitrification, ammonium transporters, mineralization of organic N and ammonium assimilation via GDH (*gudB*) and GS/GOGAT (*glnA*, *gltD*). Dissimilatory processes, often overlooked in soil N cycling studies (Moreau et al., 2019), can eventually lead to loss of gaseous nitrogen from the system either as N_2_, N_2_O or NO. As N is commonly a limiting nutrient in the rhizosphere, coveted by both plants and microbes, dissimilatory processes might be disadvantageous in this habitat. Indeed, plants have been demonstrated to outcompete nitrate reducing bacteria for nitrate (Moreau et al., 2015) and there is some evidence that root exudates can inhibit denitrification (Dassonville et al., 2011). Some of the genes involved in dissimilatory processes were also significantly upregulated in litter-amended bulk soil compared to litter-amended rhizosphere, also suggesting the inhibition of N-loss processes in rhizosphere soil. Expression of assimilatory processes that were upregulated in the detritusphere compared to bulk soil (*gudB, gltD*) was higher than that of any DNRA or denitrification genes. Assimilatory processes are possibly a more common strategy for N acquisition in the detritusphere, and their upregulation compared to unamended soil is likely driven by carbon availability from root litter to maintain C:N cellular stoichiometry. Compared to bulk soil, both detritusphere and rhizosphere have a much higher oxygen consumption due to C utilization and biomass increase. However, in the rhizosphere there is extreme competition over nitrate against the plant, whereas in the detritusphere the driving force is only C and O consumption which may create anaerobic or microaerobic niches that would support DNRA and denitrification.

Bulk soil was characterized by upregulation of genes coding for nitrification, another process that leads to loss of N from the system through generation of nitrous oxide and leaching of nitrate from soil (Prosser et al., 2020). Ammonia monooxygenase subunit C (*amoC*) was also one of the most highly expressed genes in this soil. Interestingly, the expression of subsequent steps in bacterial nitrification (*hao, nxrA, nxrB*) was extremely low. However, expression of nirK, traditionally considered a part of the denitrification process, was one of the highest we detected. When we explored the phylogeny of detected amoA and nirK variants, we discovered that all highly transcribed variants were archaeal. nirK has been detected in archaeal genomes and the current leading hypothesis is that archaea oxidize hydroxylamine by combining it with nitric oxide (NO), yielding two nitrite molecules, one of which is reduced back to NO by *nirK* (Prosser et al., 2020).

While it is possible that conditions are more favorable for nitrification in bulk soil as there is less competition over ammonium in the absence of plant roots, it is also possible that nitrification was downregulated or inhibited in the rhizosphere. Biological nitrification inhibition (BNI) by the exudation of secondary metabolites from plant roots that can suppress ammonia oxidation by nitrifying microorganisms (Subbarao et al., 2007, 2013), thus increasing the availability of ammonium to plants. Hence it is proposed as one strategy to increase crop yields and reduce fertilizer loss (Coskun et al., 2017). Root exudates of *A. fatua* have been shown to inhibit nitrification by ammonia oxidizing bacterium *Nitrosomonas europea* (O’Sullivan et al., 2017). If *A. fatua* inhibited ammonia oxidizing archaea in order to increase ammonium availability to its roots, it would likely have appeared as upregulation in bulk soil. Given that microbes in the rhizosphere appear to be experiencing N-limitation, our observations from the rhizosphere and bulk soil may be partially explained by competition over ammonium between rhizosphere microorganisms and the plant.

Despite the low relative abundance of AOA, archaeal genes for ammonia monooxygenase (*amoABC*) were among the most highly expressed out of all N-cycling transcripts in this study across time, location (bulk soil vs. rhizosphere) and presence or absence of litter amendment. The relative abundance of AOA in our study based on 16S-rRNA gene abundance, while never exceeding 1% of the microbial community, was several-fold higher than that of ammonia oxidizing bacteria (AOB) and matches previous results from other grasslands (Leininger et al., 2006; Nicol et al., 2008; Zeglin et al., 2011). While they may be very slow growers, AOA can be highly metabolically active in soil (Pratscher et al., 2011). Moreover, transcription of archaeal *amoA* has also been shown to be correlated to nitrification rates (Zhang et al., 2010; Pratscher et al., 2011; Orellana et al., 2019). A similar experiment using a different species of *Avena* demonstrated that gross nitrification rates decreased with rhizosphere age (Herman et al., 2006). This pattern further supports correlation between *amoA* expression and nitrification rates, and could reflect increasing competition over labile N as well as reduced root exudation by the plant in aging rhizosphere (Jackson et al., 1988; Hu et al., 2001; Zhalnina et al., 2018).

Expression of *amoA* has been previously shown to be dominated by few archaeal phylotypes (Tourna et al., 2008; Offre et al., 2009; Gubry-Rangin et al., 2011). Not only do our results support this, but they quantitatively show that on average 98% of the reads mapped to *amoA* were assigned to three variants, all archaeal. All three phylotypes were within the *Nitrososphaera* clade, commonly found in soil (Tolar et al., 2017). Similarly, *nirK* expression was also dominated by three variants, but surprisingly these were most closely related to another soil AOA genus – *Nitrosocosmicus*. Additionally, expression of *amoA* and *nirK* is well correlated. It is possible that Nitrososphaera are performing ammonium oxidation and *Nitrosocosimus* oxidizing the product. While there are ammonia oxidizing and nitrite oxidizing bacteria, such a cooperation is not known in archaea. However, it is also possible that the taxonomic placement of one of the genes is wrong. Both *amoA* and *nirK* have been identified in curated genomes of AOA (Jung et al., 2014; Bayer et al., 2016; Carini et al., 2018) which raises the possibility that the same AOA cell can enact both steps of nitrification (Kozlowski et al., 2016; Reji et al., 2019), but dual expression of amoA and nirK by AOA has not yet been shown experimentally. Our strong correlation suggests that there may be a relationship between these two processes, but further experimentation is required to determine if they are mechanistically linked.

Thaumarchaeal extracellular proteases identified in the same dataset were previously found to be upregulated in bulk soil <u>(Sieradzki et al. 2022)</u> and their temporal expression patterns highly correlated with *amoA* expression. Thaumarchaeota have been shown to prefer ammonium from organic nitrogen sources such as urea, amino acids and peptides in forest soil (Levičnik-Höfferle et al., 2012). As AOA have to compete with the plant over ammonium in the rhizosphere, they may turn to organic N instead (Craine et al., 2007; Blagodatskaya and Kuzyakov, 2008), explaining the tight correlation between *amoA* and AOA proteases.

Ammonium assimilation via glutamate synthase (GS/GOGAT) is generally thought to be preferred to glutamate dehydrogenase (GDH) under N limitation due to the K_m_ of GS/GOGAT which is an order of magnitude lower than that of GDH (Sakamoto et al., 1975; Alibhai and Villafranca, 1994). In our study GS/GOGAT gene expression was widespread across treatments and over time, with significant upregulation in response to both roots and litter. Moreover, the expression of GS/GOGAT was consistently at least an order of magnitude higher than that of GDH. The first step of the GS/GOGAT pathway, *glnA* (glutamine synthetase), was one of the most highly expressed genes in our study. The P_II_ regulatory protein *glnB* which is activated during N limitation and activates GS/GOGAT over GDH (Huergo et al., 2013) was also highly expressed under all experimental conditions and upregulated in the presence of live or dead roots. In contrast, expression of single-enzyme pathway glutamate dehydrogenase (GDH) was much lower. Assuming comparable downstream post-transcriptional regulation, a preference for multi-step unidirectional pathway GS/GOGAT (Hochman et al., 1988) as opposed to the bidirectional GDH, which could also perform glutamate deamination (Melo-Oliveira et al., 1996), could result from nitrogen limitation, as it prevents nitrogen loss from the organism (Yuan et al., 2009).

We identified differences in expression of GS/GOGAT and GDH between habitats. Expression of *glnA* (GS/GOGAT) was higher in the rhizosphere compared to bulk soil at 3, 6 and 12 days, and was higher in litter amended soil compared to unamended both in bulk soil and in the rhizosphere. Its expression was highest in the presence of both live and dead roots (litter-amended rhizosphere) at 3 and 6 days, indicating an additive effect of live roots and root litter. This effect may be due to high resource demand when C is available by both litter and exudates, which exacerbated the demand for N to maintain cellular stoichimetry (Zhu et al., 2014; Moreau et al., 2019). Expression of carbohydrate active enzymes (CAZy) in the same soil was also higher in the young rhizosphere and decreased over time and expression of extracellular protease genes was highest at 3 days (Nuccio et al., 2020; Sieradzki et al., 2020). This additive effect, generally accepted to be short-term (Kuzyakov et al., 2000), disappears by 6 days of rhizosphere maturation.

Temporal effects of ammonium assimilation gene expression varied by gene, pathway and habitat. For example, the two GDH genes we identified were upregulated in different habitats: rhizosphere (*gdh2*) and detritusphere (*gudB*). The expression of both genes decreased over time, possibly because ammonium concentrations dropped to a concentration too low for the affinity of this enzyme (Alibhai and Villafranca, 1994). The expression of *glnA* (GS/GOGAT) tells a more complicated story. In the litter-amended rhizosphere its expression decreases over time, as does the expression of its regulatory genes. However, in the litter-amended bulk soil the expression of *glnA* increases from day 6 on. The consistently lower expression of this enzyme in unamended bulk soil implies that any C input, be it from rhizodeposition of root litter, creates a demand for N that is at least somewhat alleviated via the GS/GOGAT pathway. Finally, we posited that there is a trade-off between the GDH and GS/GOGAT pathways that corresponds to nitrogen limitation. Therefore, we examined the ratio of gene expression between *glnA* (GS/GOGAT) and the more highly expressed of the GDH genes, *gudB*. This ratio was consistently higher than 10, implying a general preference for GS/GOGAT, possibly due to low N availability at all times with or without the presence of roots or root litter. This ratio also increased over time under all experimental conditions, but this increase began earlier in the litter-amended rhizosphere. However, by 22 days the highest ratio was in litter-amended bulk soil, possibly because growth rates are slower in bulk soil compared to the rhizosphere (Pett-Ridge et al., 2021) and therefore N limitation developed slower.

## Conclusions

Here we present a comprehensive analysis of expression of all nitrogen cycling pathways over time and in the main soil habitats: rhizosphere, litter-amended rhizosphere, litter-amended bulk soil and unamended bulk soil. We propose several classes of controllers on the expression of nitrogen cycling genes in soil with or without roots and/or root litter. Processes that lead to loss of N from the system were downregulated near live roots. Availability of carbon (C) and the quality of that C, whether from root exudates or root litter, could drive N uptake to maintain cellular stoichiometry, and proximity to live roots can lead to competition for inorganic nitrogen. The use of metatranscriptomics allowed us to track multiple pathways and identify trade-offs between them which are likely based on N availability and limitation.

## Acknowledgements

The authors would like to thank Dr. Graeme Nicol, Dr. Christina Hazard, Dr. Kataryna Zhalnina, Dr. Rachel Neurath, Dr. Rachel Hestrin, Dr. Nameer Baker, Alexa Nicholas, Katerina Estera-Molina, Dr. Spencer Diamond and Dr. Jillian Banfield for insightful discussions and emotional support during the covid-19 pandemic. This research was supported by the U.S. Department of Energy Office of Science, Office of Biological and Environmental Research Genomic Science program under Awards DE-SC0020163 and DE-SC0016247 to M.K.F at UC Berkeley and awards SCW1589 and SCW1678 to J.P-R. at Lawrence Livermore National Laboratory. Work conducted at Lawrence Livermore National Laboratory was supported under the auspices of the U.S. DOE under Contract DE-AC52-07NA27344. Sequencing was conducted as part of Community Sequencing Awards 1487 to J.P.R. and 1472 to M.K.F.. E.T.S was partially supported by Marie Sklodowska-Curie postdoctoral fellowship “DIVOBIS”.

