## Supplementary figures and images for "Rhizosphere and detritusphere habitats modulate expression of soil N-cycling genes during plant development"

### Supplemental figures

#### Supplementary figure 1

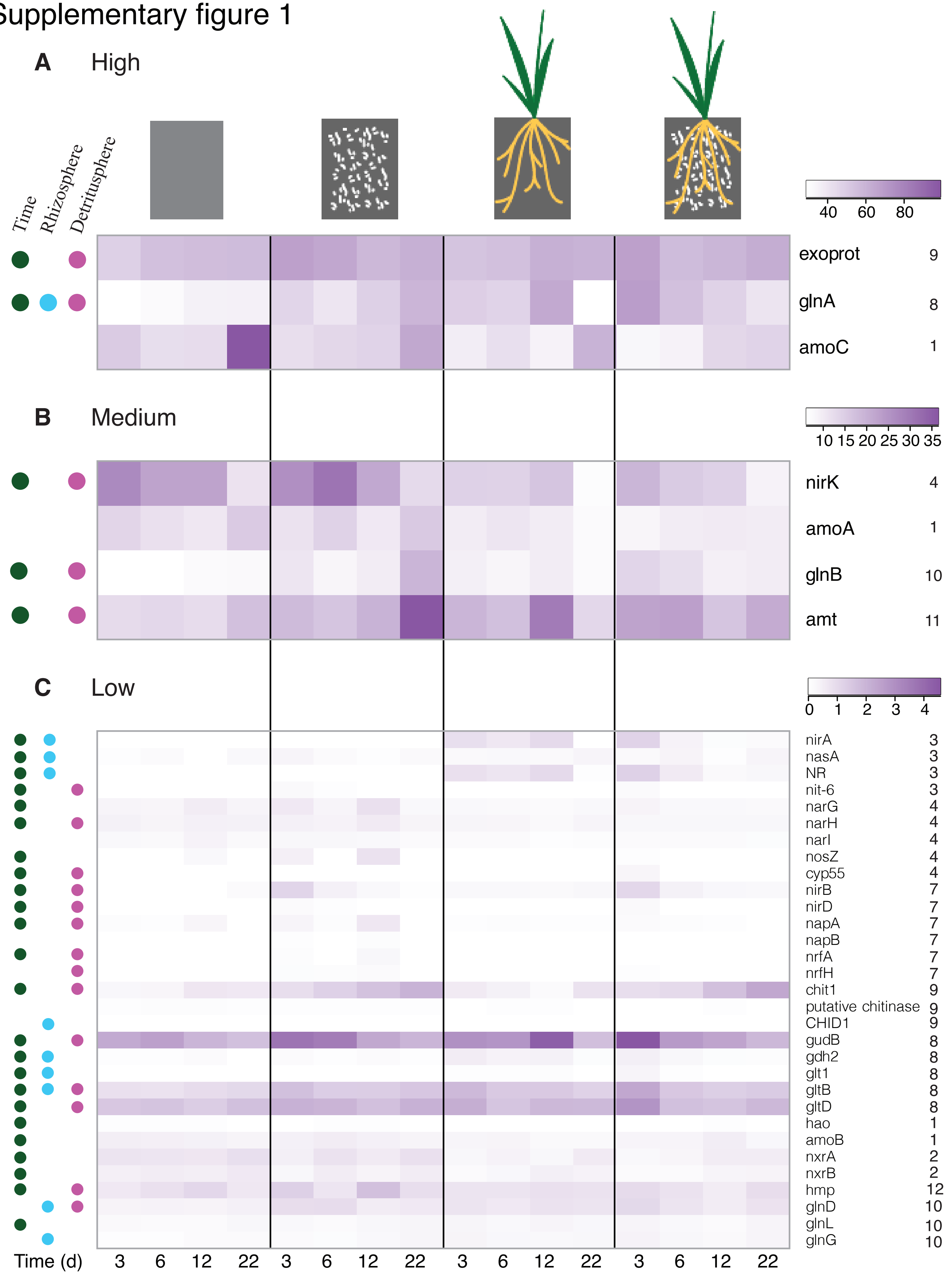

Supplementary figure 2

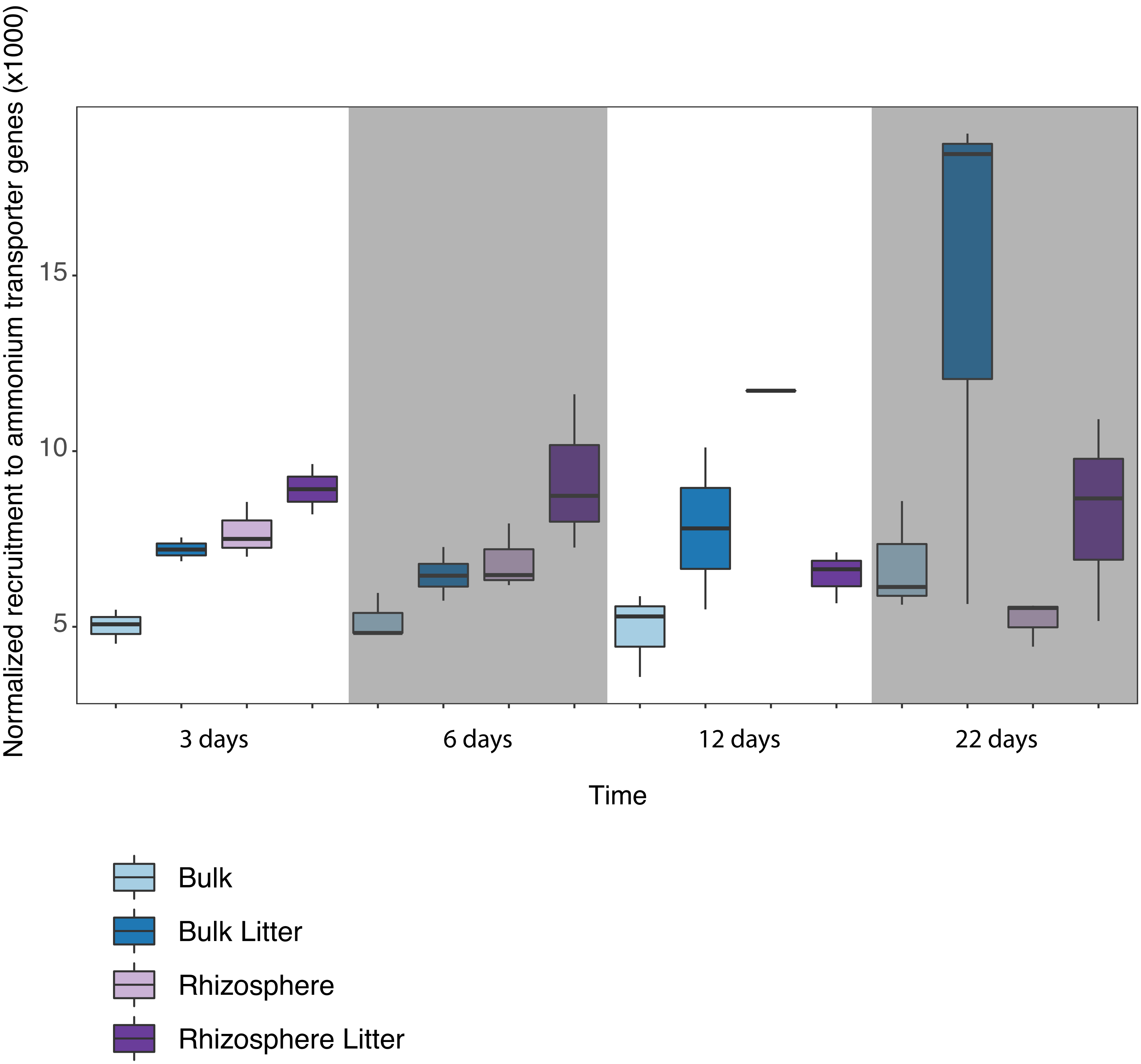

#### Supplementary figure 3

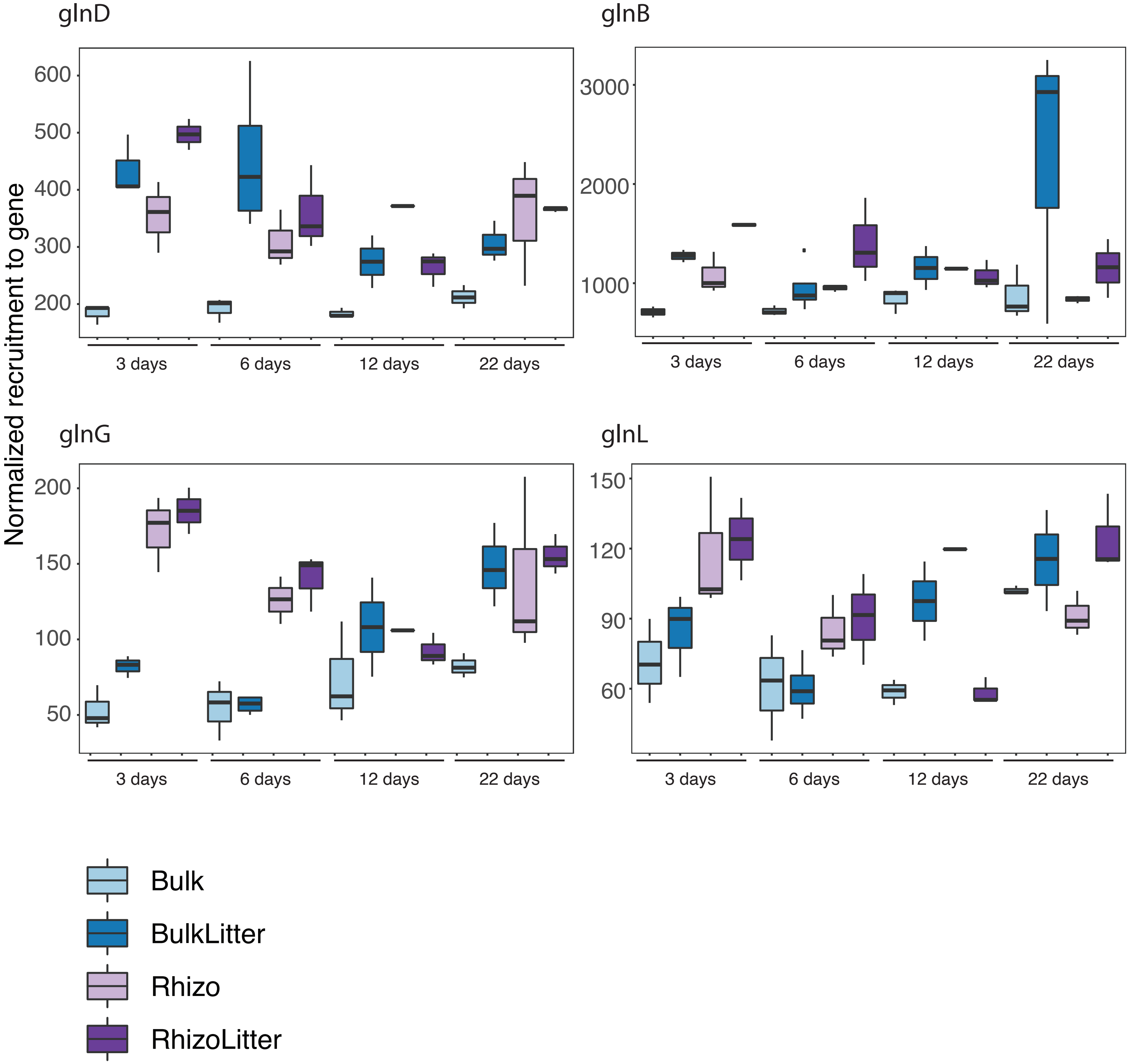

Supplementary figure 4

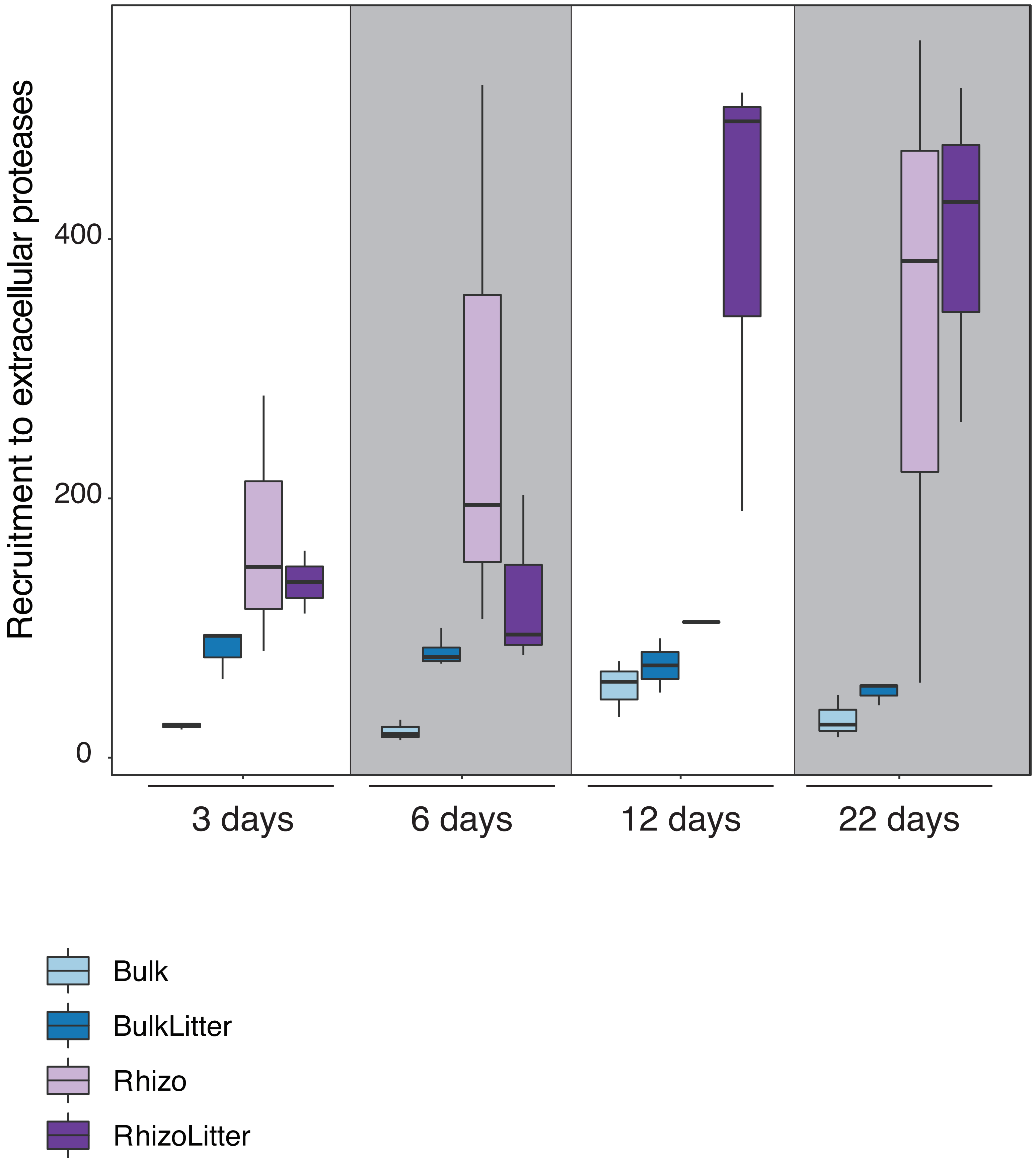
